# The Microtubule Severing Protein UNC-45A Counteracts the Microtubule Straightening Effects of Taxol

**DOI:** 10.1101/2023.09.12.557417

**Authors:** Asumi Hoshino, Valentino Clemente, Mihir Shetty, Brian Castle, David Odde, Martina Bazzaro

## Abstract

UNC-45A is the only known ATP-independent microtubule (MT) severing protein. Thus, it severs MTs via a novel mechanism. *In vitro* and in cells UNC-45A-mediated MT severing is preceded by the appearance of MT bends. While MTs are stiff biological polymers, in cells, they often curve, and the result of this curving can be breaking off. The contribution of MT severing proteins on MT lattice curvature is largely undefined. Here we show that UNC-45A curves MTs. Using *in vitro* biophysical reconstitution and TIRF microscopy analysis, we show that UNC-45A is enriched in the areas where MTs are curved versus the areas where MTs are straight. In cells, we show that UNC-45A overexpression increases MT curvature and its depletion has the opposite effect. We also show that this effect occurs is independent of actomyosin contractility. Lastly, we show for the first time that in cells, Paclitaxel straightens MTs, and that UNC-45A can counteracts the MT straightening effects of the drug. **Significance:** Our findings reveal for the first time that UNC-45A increases MT curvature. This hints that UNC-45A-mediated MT severing could be due to the worsening of MT curvature and provide a mechanistic understanding of how this MT-severing protein may act. UNC-45A is the only MT severing protein expressed in human cancers, including paclitaxel-resistant ovarian cancer. Our finding that UNC-45A counteracts the paclitaxel-straightening effects of MTs in cells suggests an additional mechanism through which cancer cells escape drug treatment.

## Introduction

MTs are stiff biological polymers [1, 2], yet, they often curve, and the result of this curving can be breaking [3, 4]. MT curvature is regulated by mechanical interactions with other cellular filaments, cell membranes, and cellular organelles, exposure to MT targeting agents, and interaction with Microtubule-Associated-Proteins (MAPs). Regarding the effect of MT targeting agents on MT curvature, *in vitro* studies on the structure of Taxol-MT have demonstrated its impact on both individual protofilaments and the overall lattice of MT. Taxol straightens the protofilaments, while simultaneously enhancing heterogeneity and flexibility within the MT lattice [5, 6]. With regards to the effect of MAPs on MT curvature, *in vitro* studies have demonstrated that the MT plus-end destabilizing proteins kinesin-13 and kinesin-1 stabilize the *αβ*-tubulin curved conformation and lock the MT lattice in a curved conformation, respectively [7] [8], while the MT stabilizing protein Tau distorts the *αβ*-tubulin within MT filaments and increases MT curvature [9]. The contribution of MT severing proteins on MT curvature has never been described.

The uncoordinated protein 45 (UNC-45) plays a significant role in various organisms due to its wide distribution and evolutionary conservation [10, 11]. In invertebrates, there exists a single isoform of UNC-45 that regulates the stability and functioning of myosin. Vertebrates, on the other hand, have two isoforms of UNC45: UNC45B, found exclusively in muscle cells, and UNC45A, expressed in all cell types [12]. Despite its conservation throughout evolution, the precise functions of UNC-45 are not yet fully understood. UNC-45A comprises four distinct domains, namely an N-terminal domain that contains three tetratricopeptide repeat (TPR) sequences. We recently made the discovery that the N-terminal domain of UNC-45A is essential for microtubule MT binding [13]. Additionally, there is a central domain whose function remains largely unknown, a neck domain that has recently been proposed as necessary for the oligomerization of UNC-45 [14], and a C-terminal UCS domain that plays a critical role in interacting with myosin II [15, 16].

In mammalian cells UNC-45A exhibits a dual role that is not mutually exclusive, functioning as a regulator of both myosin and MT activities. With regards to the latter, UNC-45A is a centrosome-[17] [18] and mitotic-spindle associated protein in cancer cells [19] [17], and is enriched in MT dense areas of nervous system and ciliated epithelium [20]. UNC-45A exhibits upregulated expression in cancer cells, wherein it antagonizes the MT-stabilizing properties of Paclitaxel, enabling cells to undergo division via multipolar spindles despite the presence of the drug. Conversely, the inhibition of UNC-45A reestablishes the susceptibility of cancer cells to Paclitaxel, thereby restoring their sensitivity to the drug [19]. Mechanistically, our recent findings have shown that UNC-45A is a MT severing protein in both *in vitro* and in living cells and that this effect is independent of myosin [21–24]. What makes this particularly intriguing is that, unlike all other known MT severing proteins, UNC-45A lacks an ATPase domain and affects MT stability independent of ATP [13].

Because we have observed the appearance of MT bends and kinks prior to UNC-45A-mediated MT severing in both *in vitro* and living cell experiments [13], here we wanted to ascertain the position of UNC-45A with respect to MT curvature and determine if UNC-45A expression affects MT curvature in cells. We found that UNC-45A enriched in the areas where MTs are curved versus the areas where MT is straight. We also show that UNC-45A overexpression increases the curvature of both perinuclear and peripheral MTs. This effect of UNC-45A on MT curvature is independent of actomyosin contractility because UNC-45A binds to MTs independent of its myosin II binding domain and acts on MT even in the presence of the myosin II inhibitor blebbistatin [13, 19]. Furthermore, here we show that actomyosin contractility does not affect perinuclear MT curvature. Lastly, we show that Paclitaxel straightens MTs in cells and that UNC-45A counteracts the MT straightening effects of Paclitaxel. Taken together, our studies support the role of the MT severing protein UNC-45A as a regulator of MT curvature in cells including in cells exposed to the MT-targeting agent Paclitaxel.

## Results

### UNC-45A signal increases in the curved region of MTs, and in Taxol-stabilized GTP MTs

We have recently shown exposure to UNC-45A kinks *in vitro* MTs and that MTs’ break and depolymerization follow these kinks [13]. We have also demonstrated that MTs in UNC-45A overexpressing cells (mouse rat fibroblasts, RFL-6) break and these breakages are preceded by MT bending at the breaking site [13]. While performing these experiments, we noticed that MTs in UNC-45A overexpressing cells appeared more curved overall than control cells. Thus, we sought to determine the localization of UNC-45A with respect to MT curvature. For this, we used the *in vitro* system because it allows for the evaluation of UNC-45A with respect to MT curvature in the absence of other cellular factors and MTs with a broad range of curvatures and because it allows evaluating single MTs in the absence of the MTs crowdedness typical of cells. GMPCPP-stabilized and rhodamine-labeled MTs were further stabilized with 10 μM Taxol as we have previously described [13, 19], and gently pipetted in a perfusion chamber so that a subset of the MTs would curve naturally while binding to the surface of the chamber. Next, we introduced 250 nM of GFP-UNC-45A and imaged the localization of UNC-45A to MT curvature using TIRF microscopy as we have previously described [13, 19] (Figure 1A). As shown in Figure 1B-C, we found that UNC-45A is enriched in the areas where MTs are curved versus the areas where MT is straight. To provide additional evidence of UNC-45A’s preference for curved MTs, we conducted a comparative analysis of GFP-UNC-45A signal intensity between two types of *in vitro* MTs: rhodamine-labeled GMPCPP MTs and Taxol-stabilized GTP MTs. Previous studies have demonstrated that these two MT types exhibit distinct characteristics in terms of stiffness and curvature with rhodamine-labeled GMPCPP MTs being more rigid and predominantly straight, and Taxol-stabilized GTP MTs being less rigid and more curved [1, 25]. We found that UNC-45A signal is enriched in the Taxol-stabilized GTP MTs versus the GMPCPP MTs (Figure 1D-M). Taken together, this suggests that UNC-45A preferentially binds to curved MTs.

**Figure 1.**
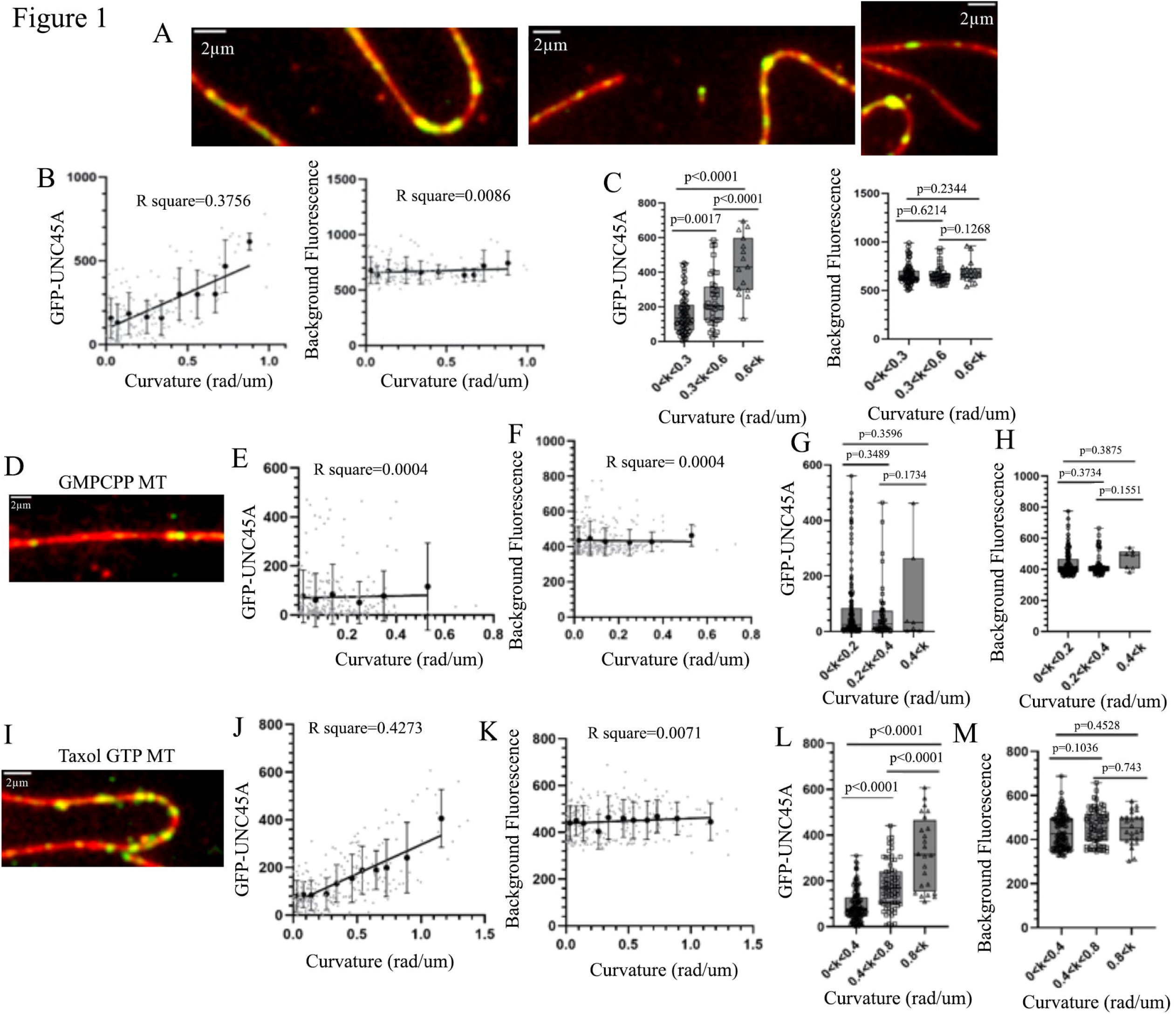
UNC-45A is enriched in the curved regions of MTs and in Taxol stabilized GTP MTs. **A**. Representative images of GFP-UNC-45A (*green*) binding to straight and curved rhodamine-labeled Taxol-stabilized GMPCPP MTs (*red*). **B**. *Left*, GFP-UNC-45A signal intensity (expressed as Arbitrary Fluorescence Intensity-AFU). *Right*, background signal intensity of green channel (expressed as AFU) nearby MT curves plotted against MT curvature. Sixteen MTs were analyzed for a total of 137 measurements from 3 different chambers. Single measurements are shown as gray dots, and average intensity and standard deviations are shown as black dots with error bars. **C**. *Left*, GFP-UNC-45A signal intensity (expressed as Arbitrary Fluorescence Intensity-AFU) categorized into three MT curvature k groups (0<k<0.3 rad/µm, 0.3≤k<0.6 rad/µm, and 0.6≤k rad/µm). *Right*, background fluorescence intensity of green channel (expressed as AFU) nearby MT curves are categorized into three MT curvature k groups (0<k<0.3 rad/µm, 0.3≤k<0.6 rad/µm, and 0.6≤k rad/µm). **D**. Representative images of GFP-UNC-45A (*green*) binding to rhodamine-labeled GMPCPP MT (*red*). **E**. GFP-UNC-45A signal intensity (expressed as Arbitrary Fluorescence Intensity-AFU). **F.** Background signal intensity of green channel (expressed as AFU) nearby MT curves plotted against GMPCPP MT curvature. Twenty MTs were analyzed for a total of 245 measurements from 3 different chambers. **G**. GFP-UNC-45A signal intensity (expressed as Arbitrary Fluorescence Intensity-AFU) categorized into three MT curvature k groups (0<k<0.2 rad/µm, 0.2≤k<0.4 rad/µm, and 0.4≤k rad/µm). **H.** Background fluorescence intensity of green channel (expressed as AFU) nearby MT curves are categorized into three MT curvature k groups (0<k<0.2 rad/µm, 0.2≤k<0.4 rad/µm, and 0.4≤k rad/µm). **I**. Representative images of GFP-UNC-45A (*green*) binding to rhodamine-labeled Taxol stabilized GTP MT (*red*). **J**. GFP-UNC-45A signal intensity (expressed as Arbitrary Fluorescence Intensity-AFU). **K**. Background signal intensity of green channel (expressed as AFU) nearby MT curves plotted against Taxol stabilized GTP MT curvature. **L**. GFP-UNC-45A signal intensity (expressed as Arbitrary Fluorescence Intensity-AFU) categorized into three MT curvature k groups (0<k<0.4 rad/µm, 0.4≤k<0.8 rad/µm, and 0.8≤k rad/µm). **M**. Background fluorescence intensity of green channel (expressed as AFU) nearby MT curves are categorized into three MT curvature k groups (0<k<0.4 rad/µm, 0.4≤k<0.8 rad/µm, and 0.8≤k rad/µm).

### UNC-45A overexpression increases MT curvature in the peripheral area of the cells

We investigated the influence of UNC-45A on MT curvature in the peripheral region of the cells. This area is commonly examined when studying morphological alterations in MTs, as it offers convenient visualization, enables the selection of individual MTs, and facilitates their analysis [3]. To this end, RFL-6 cells were lentivirally infected with either GFP (control) or UNC-45A-GFP and twelve hours post infection cells were fixed, stained with anti-alpha-tubulin to visualize MTs (Figure 2A) and subjected to immunofluorescence microscopy. For each condition (cells expressing GFP and UNC45A-GFP), we assessed MT curvature in visually extracted single MTs using a semi-automated MT tracking algorithm in MATLAB. This method was previously described in our work [3, 26, 27]. We focused on individual MTs that were neither crossing nor bundling and were at least 2 µm away from the cell edge. To quantify the curvature, we selected five points with the highest curvature within each condition. Using these parameters and analysis method, we found that MTs in UNC-45A-GFP overexpressing cells were more curved than control GFP overexpressing cells (Figure 2B). All measurements were taken in MTs with similar fluorescent intensity (Figure 2C, *left*) and in cells expressing equal amounts of GFP (Figure 2C, *right* and Supporting Figure 1A).

**Figure 2.**
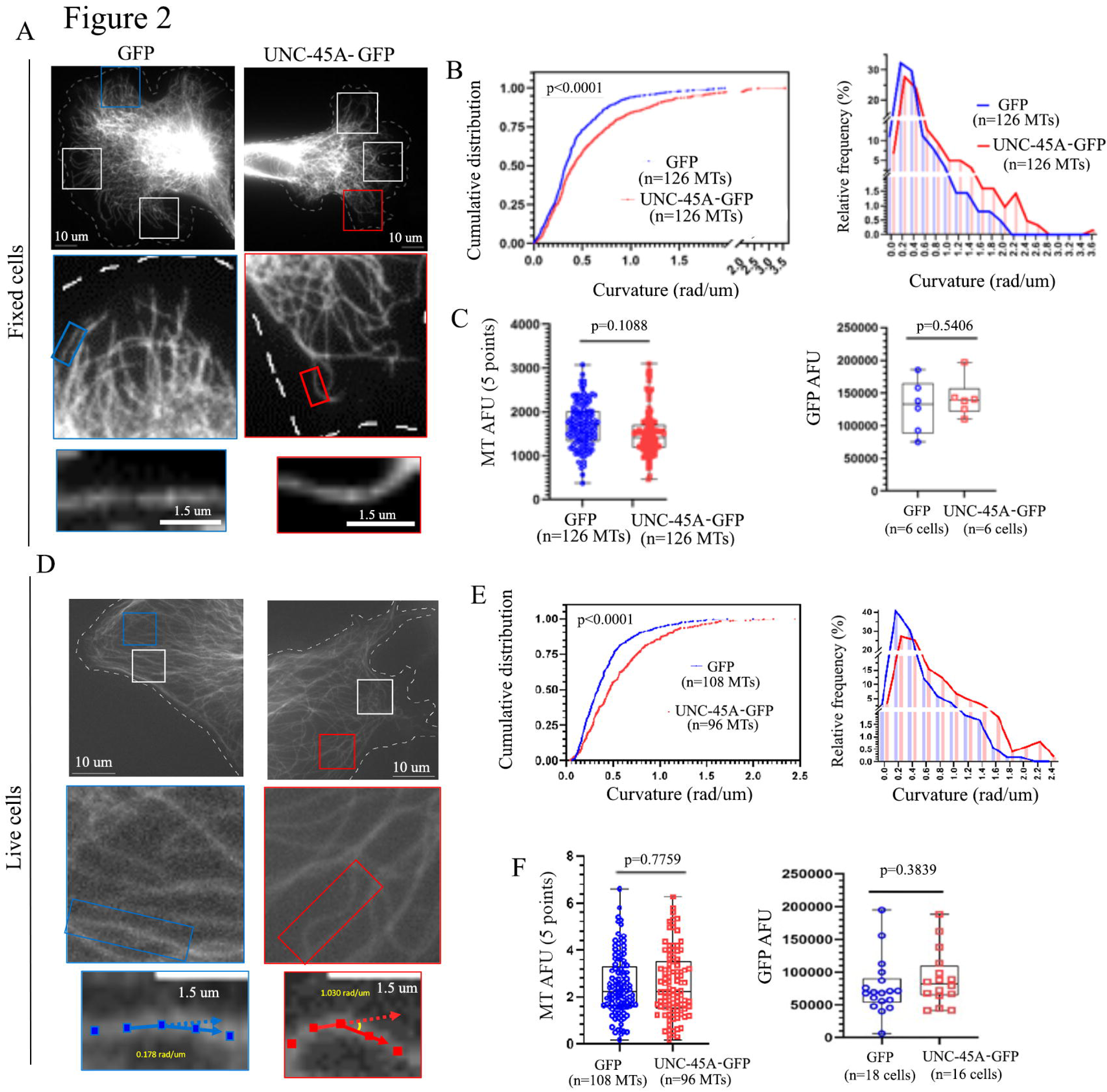
Overexpression of UNC-45A increases MT curvature in the peripheral area of cells. **A**. *Top two panels,* representative images of MTs visualized via anti-alpha-tubulin staining in fixed GFP and UNC-45A-GFP overexpressing RFL-6 cells. The white dotted lines indicate cell edges. Images were taken with the same exposure time. Three representative regions of interest (ROI) are shown. *Middle two panels*, close-up of one of representative areas shown in the top panels. *Bottom two panels*, representative magnified images region of single MT shown in middle panels where 5 curvature values were obtained. **B.** *Left,* cumulative distribution of MT curvature calculated using the 5 curvature points per condition. The mean curvature values and standard deviations of GFP and UNC-45A-GFP were 0.421 rad/*µ*m ± 0.341 and 0.581 rad/ *µ*m ± 0.509 respectively. *Right,* histogram of MT curvature distribution shown in B, left. **C**. *Left,* quantification of MT fluorescence intensity (arbitrary fluorescence units – AFU) along the length of the measured MTs. n= number of total MTs evaluated per condition. *Right,* quantification of GFP mass (arbitrary fluorescence units-AFU) in GFP and UNC-45A-GFP overexpressing cells.**D**. *Top two panels,* representative images of MTs visualized with DeepRed in live GFP and UNC-45A-GFP overexpressing RFL-6 cells. The white dotted lines indicate cell edges. Images were taken using the same exposure time. Two representative regions of interest (ROI) are shown. *Middle two panels*, close-up of one of representative areas shown in the top panels. *Bottom two panels*, representative magnified images region of single MT shown in middle panels where 5 curvature values were obtained. For each MT, dots were placed every 0.5µm for 2.5µm long to record x-y coordinates. Yellow numbers indicate calculated curvature values at middle point using two adjacent points along MT. **E**. *Left,* cumulative distribution of MT curvature calculated using the 5 curvature points per condition. The mean curvature values and standard deviations of GFP and UNC-45A-GFP were 0.415 rad/*µ*m ± 0.303 and 0.571 rad/ *µ*m ± 0.407 respectively. *Right,* histogram of MT curvature distribution shown in E, left. **F**. *Left,* quantification of MT fluorescence intensity (arbitrary fluorescence units – AFU) along the length of the measured MTs. n= number of total MTs evaluated per condition. *Right,* quantification of GFP mass (arbitrary fluorescence units-AFU) in GFP and UNC-45A-GFP overexpressing cells.

Next, our aim was to verify whether UNC-45A overexpression in living cells also results in an increase in MT curvature, similar to what we observed in fixed cells (Figure 2A-C). However, the semi-automated MT tracking algorithm we employed for fixed cells is not applicable to live cell imaging, especially when using fluorescent microscopy [3, 26, 27], which is essential for our experiments. Therefore, our initial goal was to confirm that the manual quantification of MT curvature using the click-point method, as utilized in our previous work [3, 26, 27], yields consistent results with the semi-automated approach. As depicted in Supporting Figure 1B-D, both automated and manual quantification of MT curvature in GFP and UNC-45A-GFP overexpressing cells yielded comparable results in fixed cell samples. Thus, we performed time-lapse microscopy of Tubulin Tracker Deep Red-labeled MTs in RFL-6 cells lentivirally infected with either GFP (control) or UNC-45A-GFP (Figure 2D). We found that UNC-45A-GFP overexpressing cells had MTs that were significantly more curved than in control cells (Figure 2E). We took all measurements in MTs with similar fluorescent intensity (Figure 2F, *left panel*) and in cells expressing equal amounts of GFP (Figure 2F, *right panel* and Supporting Figure 1E). Taken together, this suggests UNC-45A may act on MT curvature in cells.

### UNC-45A overexpression increases MT curvature in the perinuclear area of the cells

In the above-described set of experiments, we evaluated MT curvature in GFP, and UNC-45A-GFP expressing cells in MTs at the cell periphery. This is commonly done because it allows easy visualization, selection and analysis of individual MTs [3]. However, this cell area is subjected to intense actomyosin contractility, which can affect MT curvature [28]. To demonstrate UNC-45A’s influence MT curvature independently of actomyosin, we next focused on examining its effect on a distinct subset of cellular MTs. Specifically, we investigated the MTs located in the perinuclear region of cells. This region was chosen because previous research has suggested that it experiences fewer actomyosin-generated forces in certain cell types [28]. By concentrating on this area, we aimed to establish a clearer understanding of UNC-45A’s role in shaping MT curvature, free from potential confounding influences of actomyosin dynamics. For this, we first needed to firmly establish that that actomyosin contractility does not affect the curvature of perinuclear MTs in RFL-6 cells, the cell type that we are using in this study. RFL-6 cells were either mock-treated or treated with increasing concentrations (0.5-5 nM) of the actomyosin activator calyculin A [29]. After 10 minutes of calyculin exposure cell lysates were subjected to Western blot analysis. As shown in Supporting Figure 2A, calyculin A treatment (1 and 5 nM) increased the levels of pS19 MLC as compared to the control, while it did not affect the total levels of MLC. Thus, both these concentrations activate actomyosin contractility RFL-6 cells. Next, we determined where activation of actomyosin contractility in the calyculin A exposed RFL-6 cells takes place. As shown in Supporting Figure 2B and C, immunofluorescence staining of phosphorylated MLC (phospho-Ser19-MLC) in control and calyculin A treated cells revealed a distribution of phosphorylated MLC in both perinuclear and peripheral areas of the cells with what appears to be a predominant staining in the peripheral areas. This is consistent with what is founds in human fibroblast (NIH 3T3) cells which, following calyculin A treatment, revealed higher concentrations of phospho-MLC at the cell’s periphery [30]. Having established that calyculin A treatment activates actomyosin contractility in RFL-6, we determined whether this results in changes in perinuclear MT curvature. RFL-6 cells were either mock-treated (control) or treated with 1nM of calyculin A for 10 minutes, after which they were fixed, stained with anti-alpha-tubulin to visualize MTs (Supporting Figure 2D) and subjected to immunofluorescence microscopy. Per each condition (control and calyculin A) MT curvature was evaluated as for the experiments presented in Figures 2 and 3. We found that calyculin A treatment does not affect perinuclear MT curvature (Supporting Figure 2E). All measurements were taken in MTs with similar fluorescent intensity (Supporting Figure 2F). We then performed a complementary experiment where perinuclear MT curvature was evaluated in RFL-6 cells treated with the myosin II inhibitor blebbistatin. Here, RFL-6 cells were mock treated (control) or treated with 25 μM of blebbistin for 1 hour, fixed and stained with anti-alpha-tubulin to visualize MTs (Supporting Figure 2G) and subjected to immunofluorescence microscopy. Per each condition (control and blebbistatin), MT curvature was evaluated as for the experiments presented in Figures 2 and 3 and described above. We found that blebbistatin treatment does not affect perinuclear MT curvature (Supporting Figure 2H). All measurements were taken in MTs with similar fluorescent intensity (Supporting Figure 2I).

**Figure 3:**
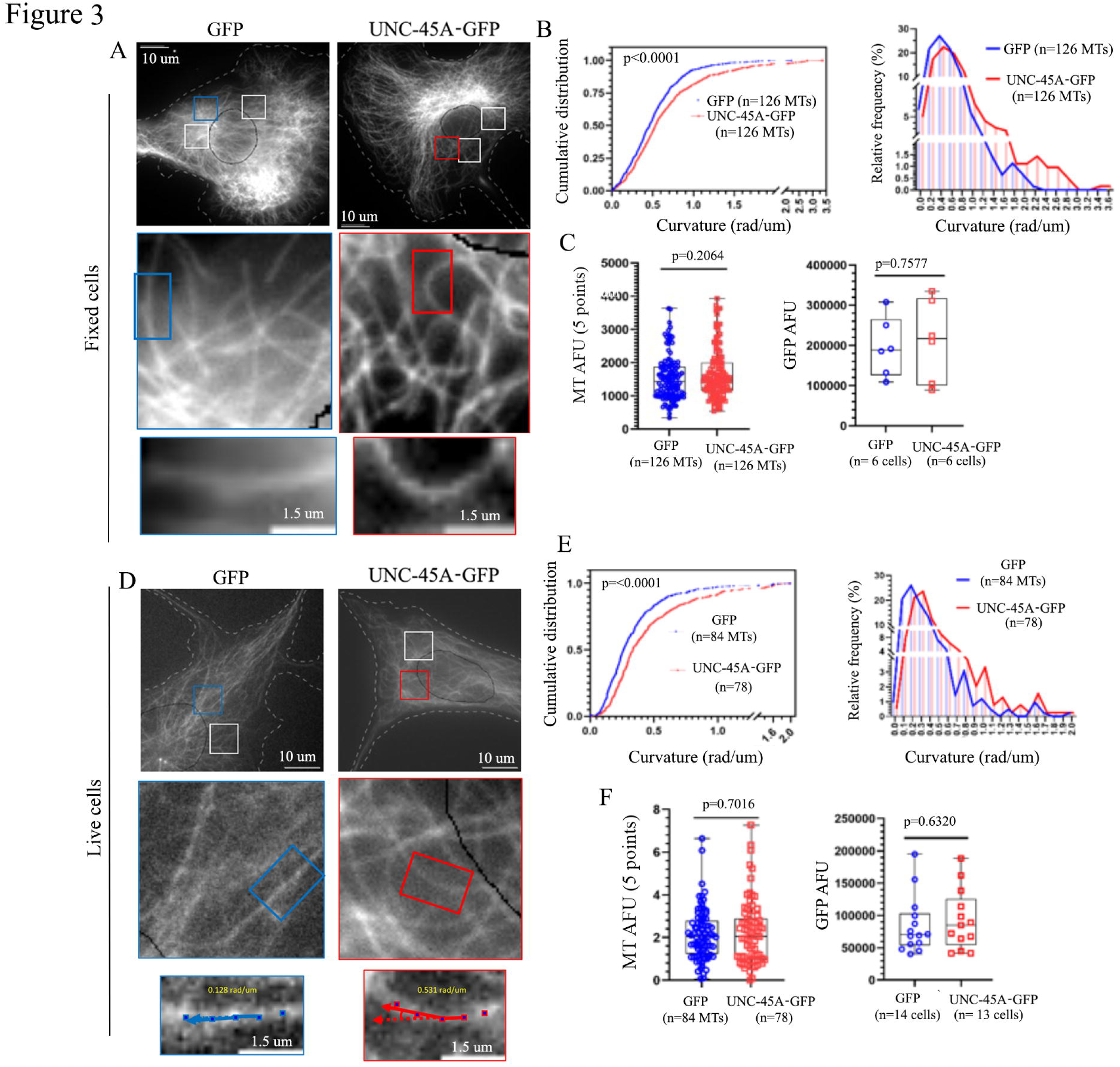
Overexpression of UNC-45A increases MT curvature in the perinuclear area of cells. **A**. *Top two panels,* representative images of perinuclear MTs visualized via anti-alpha-tubulin staining in fixed GFP and UNC-45A-GFP overexpressing RFL-6 cells. The white dotted lines indicate cell edges and black lines indicate nuclear membrane. Images were taken with the same exposure time. Three representative regions of interest (ROI) are shown. *Middle two panels*, close-up of one of representative areas shown in the top panels. *Bottom two panels*, representative magnified images region of single MT shown in middle panels where 5 curvature values were obtained. **B.** *Left,* cumulative distribution of MT curvature calculated using the 5 curvature points per condition. The mean curvature values and standard deviations of GFP and UNC-45A-GFP were 0.510 rad/µm ± 0.339 and 0.664 rad/ µm ± 0.508 respectively. *Right,* histogram of MT curvature distribution shown in B, *left*. **C**. *Left,* quantification of MT fluorescence intensity (arbitrary fluorescence units – AFU) along the length of the measured MTs. n= number of total MTs evaluated per condition. *Right,* quantification of GFP mass (arbitrary fluorescence units-AFU) in GFP and UNC-45A-GFP overexpressing cells. **D**. *Top two panels,* representative images of MTs visualized with DeepRed in live GFP and UNC-45A-GFP overexpressing RFL-6 cells. The white dotted lines indicate cell edges and black lines indicate nuclear membrane. Images were taken using the same exposure time. Two representative regions of interest (ROI) are shown. *Middle two panels*, close-up of one of representative areas shown in the top panels. *Bottom two panels*, representative magnified images region of single MT shown in middle panels where 5 curvature values were obtained. For each MT, dots were placed every 0.5µm for 2.5µm long to record x-y coordinates. Yellow numbers indicate calculated curvature values at middle point using two adjacent points along MT. **E**. *Left,* cumulative distribution of MT curvature calculated using the 5 curvature points per condition. The mean curvature values and standard deviations of GFP and UNC-45A-GFP were 0.336 rad/µm ± 0.267 and 0.451 rad/ µm ± 0.341 respectively. *Right,* histogram of MT curvature distribution shown in E, *left*. **F**. *Left,* quantification of MT fluorescence intensity (arbitrary fluorescence units – AFU) along the length of the measured MTs. n= number of total MTs evaluated per condition. *Right,* quantification of GFP mass (arbitrary fluorescence units-AFU) in GFP and UNC-45A-GFP overexpressing cells.

After confirming that actomyosin contractility has no impact on perinuclear MT curvature in RFL-6 cells, our investigation proceeded to examine the influence of UNC-45A overexpression on these perinuclear MTs. This was done in both fixed and live cells looking at MTs within 10 μm from the nuclear membrane as identified via DAPI staining. In fixed cells (Figure 3A), we found that UNC-45A overexpression led to increase in perinuclear MT curvature (Figure 3B). All measurements were taken in MTs with similar fluorescent intensity (Figure 3C, *left*) and in cells expressing equal amounts of GFP (Figure 3C, *right* and Supporting Figure 1A). We also looked at the effect of UNC-45A overexpression on perinuclear MTs curvature in live cells. We found that UNC-45A-GFP overexpressing cells (Figure 3D) had perinuclear MTs that were significantly more curved than the ones in control cells (Figure 3E). All measurements were taken in MTs with similar fluorescent intensity (Figure 3F, *left*) and in cells expressing equal amounts of GFP (Figure 3F, *right* and Supporting Figure 1E).

To confirm the effect of UNC-45A on MT curvature, we performed a complementary experiment where perinuclear MT curvature was evaluated in under conditions of UNC-45A loss. To this end, UNC-45A was knocked down via lentiviral-mediated delivery of scramble of UNC-45A shRNAs as we have previously performed [19]. Because the lentiviral vector used expresses GFP, it is possible to monitor for efficiency of lentiviral infection [13]. Western blot analysis revealed the efficiency of the KD with >60% reduction in UNC-45A expression levels (Figure 4A). Next, scramble and UNC-45A KD RFL-6 cells were fixed, stained with anti-alpha-tubulin to visualize MTs (Figure 4B), and subjected to immunofluorescence microscopy. Per each condition (scramble and UNC-45A KD cell), curvature was evaluated on perinuclear MTs (MTs that were within 10 μm from the nuclear membrane) as described above. We found that MTs in UNC-45A KD cells were less curved than scramble cells (Figure 4C). All measurements were taken in MTs with similar fluorescent intensity (Figure 4D, *left*) and in cells expressing equal amounts of GFP (Figure 4D, *right* and Supporting Figure 3). Taken together, this suggests UNC-45A may act on MT curvature independent of actomyosin forces.

**Figure 4.**
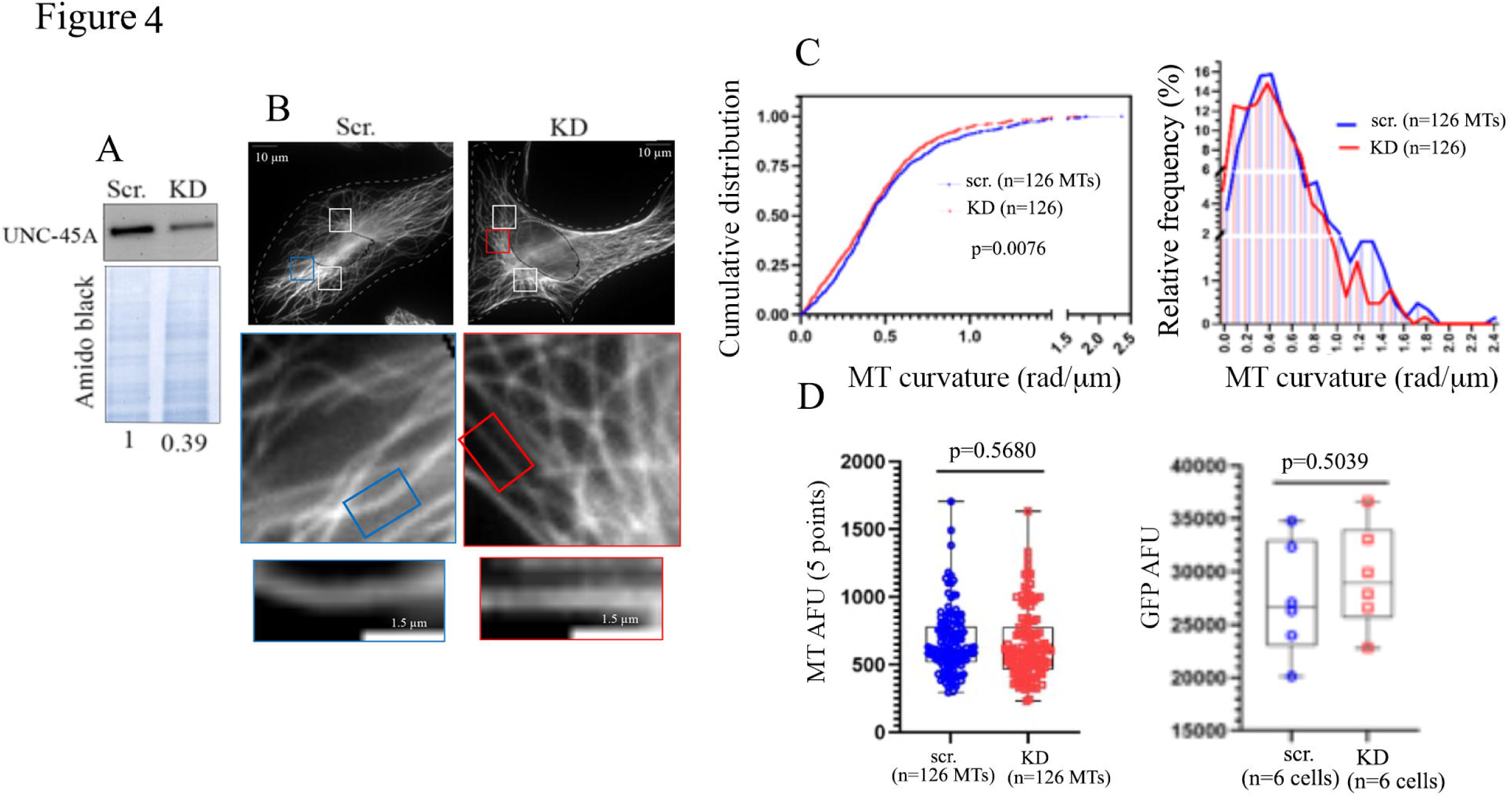
Loss of UNC-45A leads to a decrease in MT curvature. **A**. Western blot analysis of the levels of UNC-45A in scramble (Scr.) and UNC-45A knock down (KD) RFL-6 cells. Amido black was used as a loading control. Numbers represent the ratio between UNC-45A and amido black. **B.** *Top two panels,* representative images of perinuclear MTs visualized via anti-alpha-tubulin staining in fixed scramble and UNC-45A knockdown (KD) RFL-6 cells. The white dotted lines indicate cell edges and black lines indicate nuclear membrane. Images were taken using the same exposure time. Rectangles indicate the representative areas shown in the bottom panels. *Middle panels,* close up of one representative area shown in the top panels. *Bottom two panels*, magnified representative region of single MT where 5 curvature values were obtained. **C.** *Left,* cumulative distribution of MT curvature calculated using the 5 curvature points per condition. The average curvature values and standard deviations of scramble and UNC-45A KD were 0.493 rad/µm ± 0.347 and 0.444 rad/ µm ± 0.309 respectively. *Right,* histogram of MT curvature distribution shown in C, *left.* **D.** *Left*, quantification of MT fluorescence intensity (arbitrary fluorescence units-AFU) along the length of the measured MTs. n=number of total MTs evaluated per condition. *Right*, quantification of GFP mass (arbitrary fluorescence units-AFU) in scramble and UNC-45A KD cells. n=number of cells evaluated per condition.

### UNC-45A counteracts the MT straightening effects of Paclitaxel

The effect of Taxol treatment on MT curvature *in vitro* has been examined in some detail. Taxol-stabilized GTP MTs are less stiff and more wavy as compared to other *in vitro* MTs [1]. However, these Taxol treated MTs also display straighter individual tubulin protofilaments given that Taxol appears to decelerate their transition from a straight to curved conformation, contributing to this unique observation [31]. Axonemal derived MTs treated with Taxol exhibited increased flexibility compared to the untreated ones. This enhanced flexibility led to the MTs becoming straightened when exposed to a flow [25]. Addition of MT-stabilizing proteins such MAP2 and Tau to these MTs increases the rigidity of these MTs. The effect of Paclitaxel on MT curvature in cells, where MT are exposed to cellular environment including the presence of other cytoskeletal and organelle components and of cellular fluid has never been reported.

To investigate this, RFL-6 cells were either mock-treated (control) or treated with 500 nM of Paclitaxel for 30 minutes and subsequently fixed and stained with anti-alpha-tubulin antibody to visualize MTs (Figure 5A). We found that Paclitaxel treatment decreased the perinuclear MT curvature compared to the control (Figure 5B-C), indicating that Paclitaxel straightens MTs in cells. All measurements were taken in MTs with similar fluorescent intensity (Figure 5D). We have previously shown that UNC-45A is overexpressed in Paclitaxel-resistant cancer cells, where it counteracts the MT stabilizing effects of the drug leading to cancer cells’ survival [19]. In light of this and of the above-presented findings that UNC-45A increases the MT curvature, we looked at the effect of Paclitaxel on MT curvature under the condition of UNC-45A overexpression. RFL-6 cells were lentivirally infected with either GFP or UNC-45A-GFP and twelve hours post infection cells were exposed or not (control) to 500 nM of Paclitaxel for 30 min. prior fixation and immunofluorescent visualization of MT stained with anti-alpha-tubulin antibody. Per each condition (GFP *minus* or *plus* Paclitaxel-Figure 6A *left panels*, and UNC-45A-GFP *minus* or *plus* Paclitaxel-Figure 6A *right panels*) MT curvature was evaluated as we described above. We found that Paclitaxel treatment straightened MT curvature in GFP expressing cells (Figure 6B, *top panel* and 6C, *left panel*). This is consistent with our findings (Figure 5) that Paclitaxel straightens MTs in un-infected RFL-6 cells. We also found that UNC-45A-GFP expressing cells Paclitaxel treatment was still able to straighten MTs (Figure 6B, *bottom panel* and Figure 6C*, right panel*). Specifically, we found that Taxol-treated MTs curved in the presence of UNC-45A-GFP but not in its absence (Figure 6D). All measurements were taken in MTs with similar fluorescent intensity (Figure 6E) and in cells expressing equal amounts of GFP (Figure 6F and Supporting Figure 4). Taken together, this suggests that UNC-45A counteracts the MT straightening effects of Paclitaxel in RFL-6 cells.

**Figure 5:**
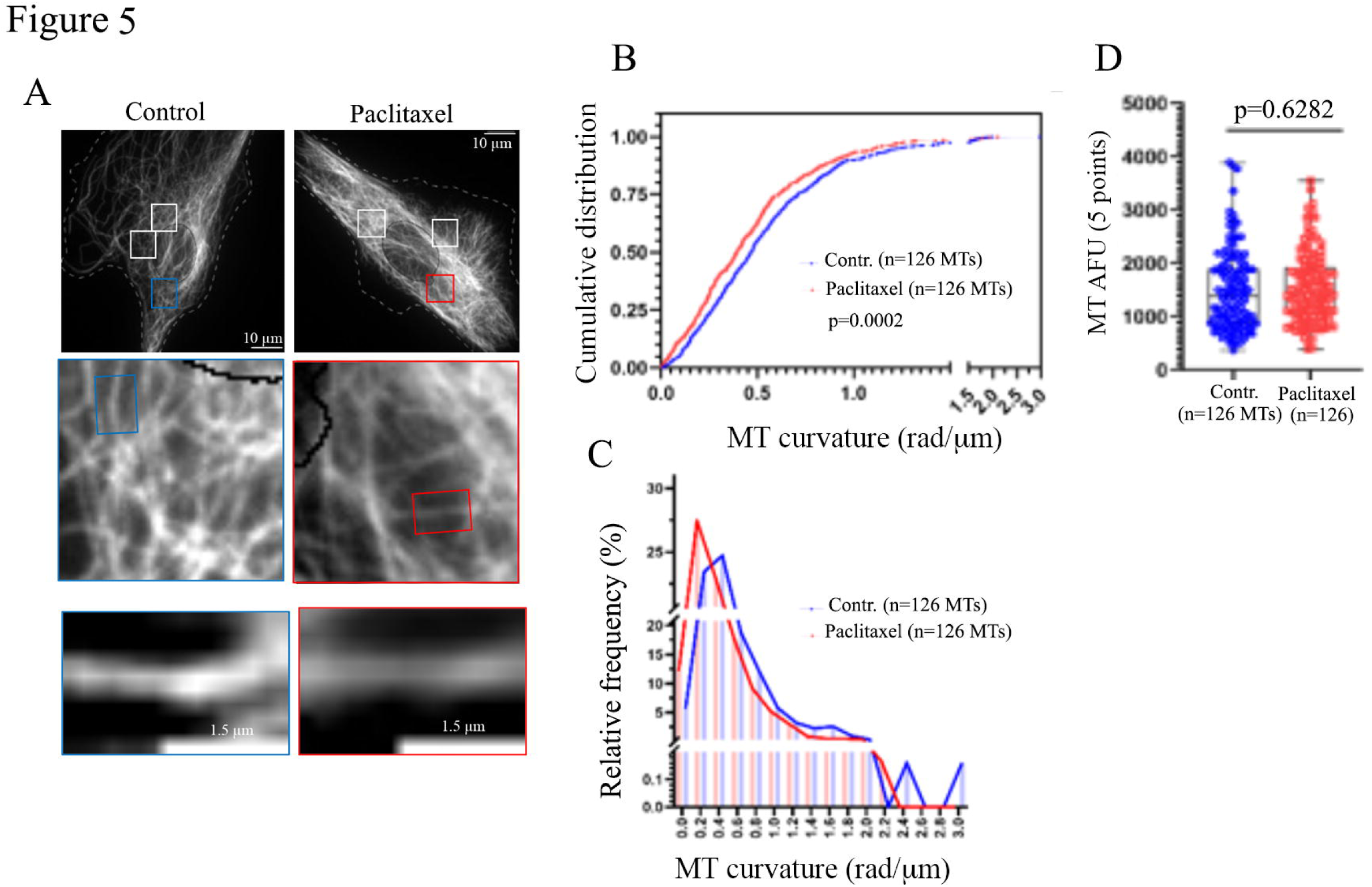
Paclitaxel straightens perinuclear MTs in cells. **A**. *Top two panels,* representative images of perinuclear MTs visualized via anti-alpha-tubulin staining in fixed RFL-6 cells treated or not (control) with Paclitaxel (500 nM for 30 min). The white dotted lines indicate cell edges and black lines indicate nuclear membrane. Images were taken with the same exposure time. Three representative region of interest (ROIs) are shown. *Middle two panels*, close-up of one of representative areas shown in the top panels. *Bottom two panels*, representative magnified images region of single MT shown in middle panels where 5 curvature values were obtained. **B**. Cumulative distribution of MT curvature calculated using the 5 curvature points per condition. The mean curvature values and standard deviations of control and Paclitaxel treatment were 0.533 rad/µm ± 0.371 and 0.455 rad/µm ± 0.343 respectively. **C.** Histogram of MT curvature distribution shown in B. **D**. Quantification of MT fluorescence intensity (arbitrary fluorescence units – AFU) along the length of the measured MTs. n= number of total MTs evaluated per condition.

**Figure 6:**
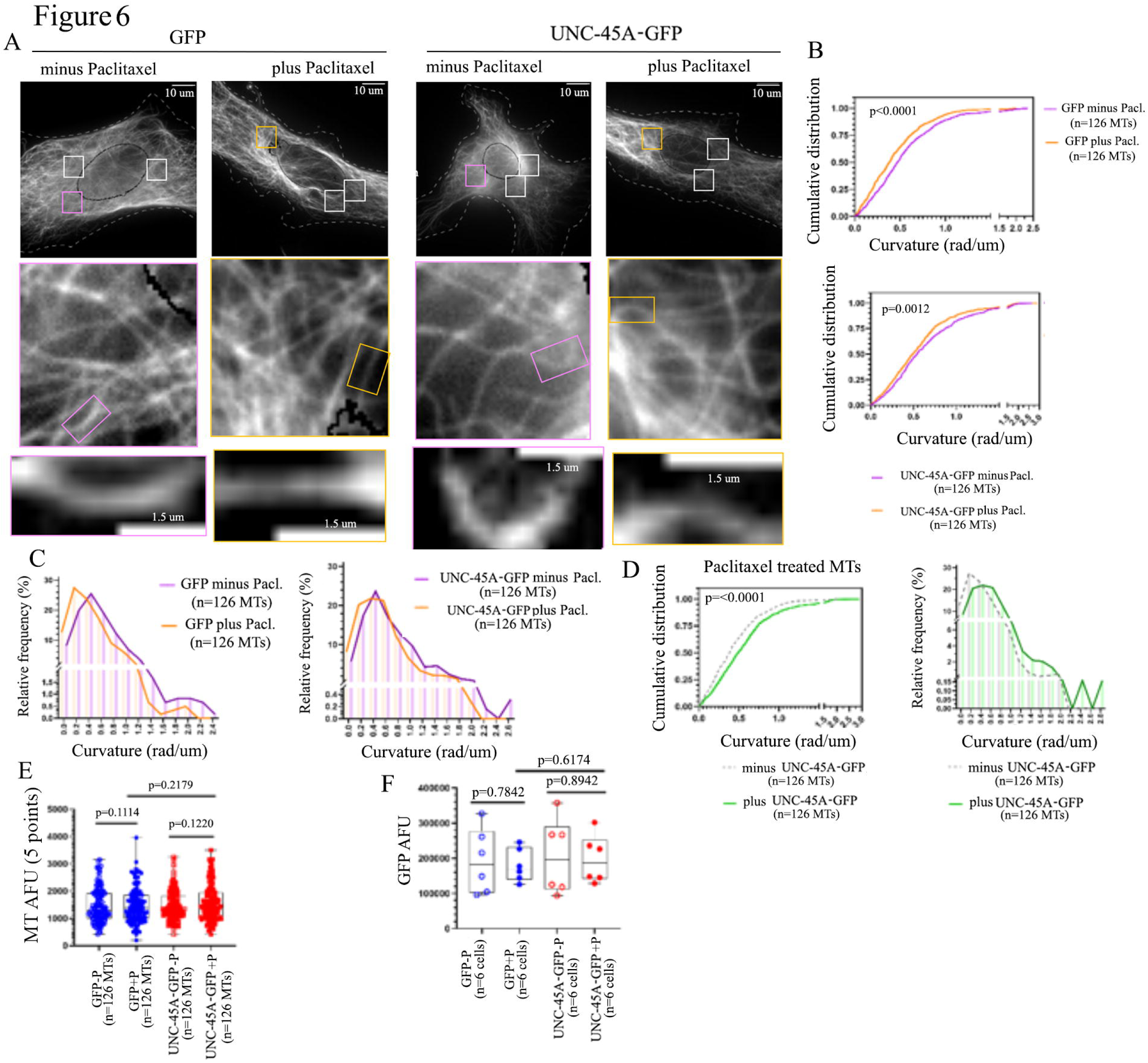
UNC-45A counteracts the MT straightening effects of Paclitaxel. **A**. *Top four panels,* representative images of perinuclear MTs visualized via anti-alpha-tubulin staining in GFP and UNC-45A-GFP overexpressing RFL-6 cells treated or not (control) with Paclitaxel (500nM for 30min) after fixation. The white dotted lines indicate cell edges and black lines indicate nuclear membrane. Images were taken with the same exposure time. Three representative regions of interest (ROI) areas are shown. *Middle four panels*, close-up of one of representative areas shown in the top panels. *Bottom four panels*, representative magnified image region of single MT shown in middle panels where 5 curvature values were obtained. **B**. *Top,* cumulative distribution of MT curvature calculated using the 5 curvature points in GFP-expressing cells per condition. The mean curvature values and standard deviations of GFP minus and plus Paclitaxel were 0.546 rad/µm ± 0.396 and 0.440 rad/ µm ± 0.328 respectively. *Bottom,* cumulative distribution of MT curvature calculated using the 5 curvature points in UNC-45A-GFP expressing cells per condition. The mean curvature values and standard deviations of UNC-45A-GFP minus and plus Paclitaxel were 0.636 rad/µm ±0.436 and 0.559 rad/ µm ± 0.403 respectively. **C**. *Left and right,* histograms of MT curvature distribution shown in B, top and bottom respectively. **D**. *Left,* cumulative distribution, *right,* histogram of Paclitaxel treated MT curvature distribution minus and plus GFP-UNC-45A. **E**. Quantification of MT fluorescence intensity (arbitrary fluorescence units – AFU) along the length of the measured MTs. n= number of total MTs evaluated per condition. **F**. Quantification of GFP mass (arbitrary fluorescence units-AFU) in GFP and UNC-45A-GFP expressing cells. n=number of cells evaluated per condition.

## Discussion

The curvature of MTs plays important roles both in normal physiological processes and in pathological conditions. Therefore, understanding the key regulators involved in MT curvature holds significance in both basic cell biology and its potential translational applications. In this study, we demonstrate that the MT severing protein UNC-45A exhibits a preferential binding to curved MTs and actively contributes to the curvature of peripheral and perinuclear MTs in cells. This binding and contribution occur independently of actomyosin contractility, as it has little to no effect on perinuclear MTs. Furthermore, our research indicates that the presence of Paclitaxel, one of the most commonly used chemotherapy agents for human cancer treatment, results in the straightening of MTs. Despite this, UNC-45A retains its ability to induce curvature in Paclitaxel-exposed MTs. This finding suggests that UNC-45A counteracts the MT-straightening effects of Paclitaxel, presenting one possible mechanism through which UNC-45A overexpression in cancer cells is associated with chemoresistance.

MTs are the stiffest of the biological polymers [1, 2], yet, curved (also referred to as bent or buckled) MTs are often found in cells in both normal and pathological [4, 32–36]. Three factors regulate MT curvature: mechanical interactions, interactions with MAPs, and exposure to MT targeting agents. Mechanical interactions MTs have with the cellular matrix surrounding them, like for instance, interaction with actin filaments at the cell cortex [3, 4, 26, 37, 38] or interaction with other cellular organelles in crowded areas of the cell, increase MT curvature [39]. MT curvature can also be influenced by their interaction with MAPs that acts on their protofilament, lattice, and/or ends, favoring or stabilizing curved or straight conformations. This is the case of kinesin-13 which depolymerizes MTs via inducing tubulin curvature and destabilization [7], of kinesin-1, which bend-locks MTs [8], or of Tau, which confers MTs’ protofilament a straight conformation [9]. Lastly, MT targeting agents can influence MT curvature. This is the case of Paclitaxel, which stabilizes straight MTs’ protofilament conformation while simultaneously increases MT flexibility *in vitro* [6, 31], or of vinblastine and colchicine, which favor and stabilize tubulin assembly and conformation, respectively [40, 41]. MT bending has physiological and pathological significance. Curved MTs are usually found in mitotic spindles [32] and crowded areas of cells [39]. Pathologically, bent and wavy MTs are often found in the axonal swelling associated with neurodegenerative diseases, aging, injuries, and axonopathies in general [35].

UNC-45A is a newly characterized MAP with ATP-independent MT severing properties [13, 20, 42]. UNC-45A-mediated MT severing is preceded by the presence of kinks and bent regions in the MTs [13]. In cells, MTs often bend and break [3, 4]. Thus, we sought to determine the contribution of UNC-45A to MT curvature in cells, if any. We started by looking at the distribution of UNC-45A on areas of MTs with different curvatures. We found that the UNC-45A signal increases in the curved region of MTs, suggesting that it may act on their curvature. This is interesting because the MT stabilizing protein Tau preferentially binds to curved MTs [43] and prevents the access of katanin [44], the most well-known of MT severing protein, to its severing site. Thus, hinting at the fact that MTs curvature may be an area of particular importance as far as MT stability regulation by severing proteins. We also found that in cells, UNC-45A actively contributes to increasing MT curvature of both peripheral and perinuclear MTs. UNC-45A has a dual non-mutually exclusive role in regulating NMII activity and MT stability [13]. Thus, we cannot exclude that the contribution of UNC-45A to peripheral MT curvature could be partly due to its effect on actomyosin contractility. However, this does not seem to be the case for perinuclear MTs since this cell area is less prone to actomyosin forces [28] and given our results that neither increasing nor decreasing actomyosin contractility affects perinuclear MT curvature in RFL-6 cells. Furthermore, because UNC-45A overexpression results in loss of overall cellular MT mass over time [13], the effect of UNC-45A overexpression on MT curvature is not due to increased MT crowdedness and instead happens despite the opposite.

*In vitro,* Paclitaxel enhances MT lattice heterogeneity by changing lateral contacts between protofilaments [5, 6] and causes structural damage (nanodamages) in the lattice [45, 46]. Both phenomena are consistent with decreased flexural rigidity of these Paclitaxel-treated MTs and increased curvature [6, 31] [47]. Our *in vitro* experiments confirm that Paclitaxel-treated MTs are curvier than the untreated ones. In cells we found that Paclitaxel straightens MTs. There are several possible explanations for what appears to be an inconsistency between *in vitro* and *in vivo* Paclitaxel-treated MTs with respect to their flexibility and curvature. First, Paclitaxel-treated MTs derived from axoneme are more flexible than untreated ones but straighter when exposed to a fluid flow [25]. Thus, it’s plausible to propose that while Paclitaxel-treated MTs may display reduced resistance to bending in RFL-6 cells, they straighten out when they align with the directional intracellular flow [48]. Second, while Paclitaxel-nanodamaged MTs *in vitro* have a softer behavior if these damages are not repaired, they become stiffer if they are repaired via free tubulin availability in the *in vitro* system [47]. Because a pool of free tubulin is present in cells, it is conceivable that Paclitaxel-treated MTs are repaired, which can account for their straightening.

Lastly and most apparent, MTs in a cell-free system are not exposed to cellular MAPs, and this also likely contributes to a difference between MT behavior in the presence of Paclitaxel between *in vitro* and cellular MTs.

We also found that UNC-45A can still curve MTs in Paclitaxel treated cells. This seemingly contradicts what we propose to be a preference of UNC-45A for curved regions of the MTs. However, as mentioned above, exposure to Paclitaxel, along with exposure to oxidative agents like hydrogen peroxide, causes nanodamages in MTs *in vitro* [45, 46, 49]. Therefore, it is possible that UNC-45A preferentially binds and acts on MT defects that are present in both curved and Paclitaxel-nanodamaged areas of MTs. Notably, UNC-45A is the only MT severing protein we know of that does not and ATP-ase domain and several human diseases like cancer and neurodegenerative diseases are characterized by both a reduction of ATP levels and a highly oxidative environment [50, 51]. In this scenario, UNC-45A may have a functional advantage over other MT destabilizing proteins and have a significant role in human diseases from cancer to neurodegeneration. Overall, our study sheds light on the pivotal role of UNC-45A in regulating MT curvature and provides valuable insights into the implications of this process in both normal and pathological human contexts. These findings may pave the way for novel therapeutic strategies targeting MT dynamics to treat human diseases.

## Experimental procedure

### Preparation of microtubules (MTs)

Porcine brain tubulin (T240) and rhodamine-labeled tubulin (TL590M) were purchased from Cytoskeleton (Denver, CO). Guanosine-5’-[(α,β)-methyleno] triphosphate (GMPCPP) was purchased from Jena Bioscience (NU-405S, Jena, Gemany). Taxol was purchased from Cytoskeleton (TXD01, Denver, CO). GMPCPP MTs were prepared as previously described with [13, 52] some modifications. Briefly, 1.5mg/mL of tubulin (5:1 mixture of unlabeled and rhodamine-labeled tubulin, respectively), 1 mM GMPCPP, and 1mM MgCl_2_ were mixed in BRB80 (80mM PIPES, 1mM MgCl_2_, 1mM EGTA, pH 6.9 – pH was adjusted with KOH) and kept on ice for 5 mins, followed by incubation at 37c for 1-2 hrs or until they were within 5-30 μm length range. Post-incubation, the seeds were diluted 10x in warm BRB80 (with 10 μM Taxol when used). Taxol-stabilized GTP MTs were prepared as previously described [53] in BRB80. Briefly, 5 mg/ml of tubulin (5:1 mixture of unlabeled and rhodamine-labeled tubulin) was polymerized in the presence of 1mM GTP for 20 min. at 37c then 50 μM of Taxol was added and incubated for additional 20 min. to equilibrate the Taxol. MTs were centrifuged at 14,000g for 10 mins at 25c and resuspended in BRB80 containing 50 μM of Taxol. The volume of BRB80 was adjusted to have similar MT density as GMPCPP MTs.

### Recombinant protein

GFP-tagged UNC-45A full-length (UNC-45A WT; 1–944 aa) was cloned, expressed, and affinity purified as previously described with some modifications to optimally preserve protein activity [13, 54]. The following detailed protocol refers to a 250 ml bacterial culture and can be scaled up or down as needed. GST-GFP-UNC-45A recombinant protein is expressed in E. coli and isolated by lysing the bacterial cell pellet using 5 mL of a phosphate-buffered saline (PBS)-based solution supplemented with 10 mM magnesium chloride (MgCl2), 1 mM phenylmethylsulfonyl fluoride (PMSF), protease inhibitor cocktail, 1% of Triton X-100 and 10% of glygerol. The cell lysis is achieved using an Avestin Emulsiflex Homogenizer C3 and the lysate is treated with 0.02 mg/ml of DNase and 400 mM of NaCl, with each treatment performed at 4°C for a duration of 15 minutes. To remove cellular debris, the lysate is subjected to centrifugation at 12,000 X g for 1 hour at 4°C. The supernatant is supplemented with 5mM DTT and added to an affinity purification column containing 1.5 mL of pre-washed Glutathione Sepharose 4B beads for 45 minutes at 4°C. At the end of the incubation period the column is washed with 5 column volumes of wash buffer (20 mM Tris-HCl pH7.5, 500mM NaCl, 10 mM of MgCl_2_, 5 mM DTT, 1% of Triton X-100 and 10% glycerol) and the protein is eluted in fractions (6X) with half column volume of elution buffer (50 mM Tris-HCl, 300mM NaCl, 10 mM of MgCl_2_, 5 mM DTT, 20 mM reduced glutathione, 1% Triton X-100 and 10% glycerol, pH=8.8). Pooled fraction-containing protein (as verified by SDS-PAGE followed by Coomassie Blue staining) are subjected to thrombin digestion to remove the GST-tag at 16°C for 4.5 hours under gentile rocking. The thrombin-to-protein ratio exhibits a variability ranging from 1:10 to 1:40 unit per microgram (μg) of protein, contingent upon the specific batches of thrombin utilized. Thrombin and GST-tag are subsequently removed from the mixture by incubation with Glutathione Sepharose 4B beads and p-aminobenzodiazepine agarose at 4°C for 45 minutes under gentle rocking. Following centrifugation, an aliquot of the supernatant-containing GFP-UNC-45A recombinant protein is subjected to SDS-PAGE and Coomassie Blue staining to determine protein purity. This protocol entails a 12-hour process, during which the protein is kept in the above describe buffer at a temperature of 4°C and protected from light. Following this, the protein undergoes centrifugation on the subsequent day to eliminate potential aggregates. The resulting supernatant is then subjected to buffer exchange (BRB80 buffer) using a spin-column with a 50kDa cut-off, and the protein concentration is quantified using absorbance at 280 nanometers (A280). It is important to note that all experimental procedures are conducted within 48 hours of the buffer exchange. Within this time frame, the recombinant GFP-UNC-45A protein remains soluble and does not undergo precipitation when stored in BRB-80 buffer at 4°C.

### Construction and preparation of flow chambers for TIRF imaging

TIRF chamber was prepared and MTs were immobilized as we previously described [13, 19]. Each channel of the TIRF chamber was rinsed with BRB80 containing 1mM DTT prior to the addition GFP-UNC-45A in imaging solution (50 mM KCl, 0.5% Pluronic F127, 0.2 mg/mL casein, 1.5% glycerol, 0.1% methylcellulose 4000 cP, 20 mM glucose, 110µg/mL glucose oxidase, and 20 µg/mL catalase, 20mM DTT in BRB80) as previously described [55]. Images were taken 10 mins after addition of GFP-UNC45A. For Taxol-stabilized MTs, the imaging buffer contained 1 μM of Taxol. Images were taken 10 mins after addition of the imaging solution containing GFP-UNC-45A.

### Total internal fluorescence (TIRF) Imaging

Time-lapse fluorescent images of MTs and GFP-UNC-45A were obtained with 561 nm and 488 nm lasers generated from TIRF mode on a Zeiss Axio observer Z1 inverted microscope using 100×/1.46 NA objective lens. An oxygen scavenging system of glucose oxidase and catalase was employed to minimize photobleaching and photodamage during illumination with the laser. The standard exposure time was 20ms for both 561 nm and 488 nm lasers.

### Cell culture

Rat RFL-6 fibroblasts were purchased from the American Type Culture Collection (ATCC) and cultured in Ham’s F-12K medium (Thermo Fisher Scientific) supplemented with 20% fetal bovine serum and 1% pen-strep. Cells were authenticated in December 2021, routinely tested negative for mycoplasma and were used between passage six and eight.

### Modulation of UNC-45A expression levels in cells

For UNC-45A overexpression and knockdown, lentiviral supernatant containing either GFP empty vector control, UNC45A-GFP (in pRRL-3’GFP vector) or scramble and UNC-45A shRNAs (in pRRL-PPTsin-3’GFP vector) were prepared and used to infect RFL-6 cells as previously described [19].

### Antibody, chemicals and tubulin

Mouse monoclonal anti-UNC-45A (Enzo, ADI-SRA-1800-D, 1:1000), phospho-myosin light chain 2 (Ser 19) antibody (Cell signaling, #3671, 1:1000), and monoclonal anti-myosin clone MY-21 (Sigma, M4401, 1:4000) were used for western blot to detect recombinant UNC-45A, p-S19 MLC and total MLC respectively. Mouse monoclonal anti-alpha-tubulin (Sigma, T6074, 1:2000) and rabbit monoclonal anti-GFP (Thermo Fisher Scientific, G10362, 1:100) were used for immunofluorescence staining. Secondary antibodies used were peroxidase-linked anti-mouse IgG and peroxidase-linked anti-rabbit IgG (both Cytiva, formerly known as GE Healthcare BioSciences, NA931 and NA934; 1:5000), Alexa Fluor 594-conjugated donkey anti-mouse IgG (1:250) and FITC-conjugated goat anti-rabbit IgG (1:200) (both Jackson ImmunoResearch Laboratories, 715-585-150 and 111-095-003, respectively). Paclitaxel was purchased from Cytoskeleton. Blebbistatin and calyculin A were purchased from Sigma-Aldrich (203390 and C5552 respectively). Tubulin Tracker Deep Red was purchased from Thermo Fisher Scientific (T34077). Porcine brain tubulin (T240) and rhodamine-labeled tubulin (TL590M) were purchased from Cytoskeleton (Denver, CO). Guanosine-5’-[(α,β)-methyleno] triphosphate (GMPCPP) was purchased from Jena Bioscience (NU-405S, Jena, Germany,). Glutathione Sepharose 4B beads (17075605) were purchased from Cytiva. DNase I (10104159001) and PMSF (10837091001) were purchased from Millipore Sigma.

### Live cell imaging of MTs in RFL-6 cells

To determine the effects of UNC-45A overexpression on MT curvature in live cells, UNC45A-GFP overexpressing RFL-6 cells (16 hrs post-infection) were treated with Tubulin Tracker Deep Red (Thermo Fisher Scientific T34077) and imaged as we have previously described [13]. Briefly, GFP-control and UNC45A-GFP overexpressing RFL-6 cells were treated with Tubulin Tracker Deep Red according to the manufacturer’s recommendation. Tubulin Tracker Deep Red was diluted 1:2000. This dilution corresponds to 500 nM Paclitaxel (personal communication from the Thermo Fisher Scientific technical support team). Cells infected with GFP control and UNC-45A-GFP overexpressing lentivirus were treated with Tubulin Tracker Deep Red sequentially so that imaging was performed under the same conditions. Time-lapse fluorescent images were collected with a Zeiss Axio observer Z1 inverted microscope using a 100×/1.46 NA objective lens. Digital images were collected at 8 s intervals over a period of 8 mins using a Cy5 filter cube. Laser power and exposure time were minimized to avoid photobleaching and photodamage, and all images for experimental and control groups were taken under the same conditions.

### Fixed cell imaging of MTs in RFL-6 cells

For studies on MT curvature in fixed cells, fixation and staining was performed as we previously described [13]. Briefly, GFP-control and UNC45A-GFP expressing RFL-6 cells or drug (blebbistatin, and Paclitaxel) treated RFL-6 cells were extracted in an MT-preserving buffer (BRB80 and 4mM EGTA) with 0.5% Triton-X for 30 secs to remove free tubulin and then fixed with 0.5% glutaraldehyde for 10 mins. After fixation, cells were quenched for 7 mins with 0.1% NaBH4 and then rehydrated in PBS before blocking in AbDil (2% TBST with 2% BSA and 0.1% sodium azide) for 30 mins. After blocking, cultures were stained with anti-α-tubulin and anti-GFP primary antibody followed by Alexa Fluor 594– and FITC-conjugated secondary antibodies. Cells were then rinsed in TBST and mounted in a Prolong Gold with DAPI. For p-S19 MLC staining, calyculin A treated RFL-6 cells were fixed in 3.7% formaldehyde in PBS and permeabilized in 0.5% Triton X-100 in TBS before staining as described [30]. Images were obtained on an Axiovert 200 microscope (Zeiss) equipped with a high-resolution CCD camera. All images were obtained using identical camera, microscope, and imaging criteria, such as gain, brightness, contrast and exposure time. Digital gray values of image pixels representing arbitrary fluorescence units (AFU) were obtained using Fiji software.

### Western blot analysis

Total cellular protein (10µg) from each sample was separated by SDS-PAGE, transferred to PVDF membranes and subjected to western blot analysis using the specified antibodies. Amido Black staining was performed to confirm equal protein loading. For detection of p-S19 MLC, RFL-6 cells were lysed in lysis buffer (50 mM Tris, pH 7.4, 150 mM NaCl, 1% Nonidet P-40, 1× protease inhibitor mixture, 1× phosphatase inhibitor mixture). For detection of UNC-45A, RFL-6 cells were lysed in 1% SDS lysis buffer.

### Drug studies

To activate actomyosin contractility, RFL-6 cells were treated with calyculin A and used concentrations varying between 0.5nM and 5nM for 10 mins to determine the optimum concentration, and RFL-6 cells were treated with 1nM calyculin A for 10 mins for p-S19 MLC and alpha-tubulin staining. To inhibit actomyosin contractility, RFL-6 cells were treated with 25µM of blebbistatin for 1h as previously described [13]. For Paclitaxel treatment, RFL-6 cells were treated with either 500nM Paclitaxel or DMSO for 30 mins.

### Analysis of MT curvature in cells

All data were recorded as 16-bit images using Zen software (blue edition, Zeiss). Single clearly visible MTs were selected, and their segments (5-10 µm length) were used to extract x-y coordinates. MT regions that were crossing and bundling with other MTs were avoided.

#### Semiautomated MT tracking

In fixed cells, three representative regions of interest (ROI) were selected within each cell and seven representative individual MTs per ROI were selected. The x-y coordinates of these MTs were extracted by semiautomated MT tacking algorism implemented in MATLAB as we have previously described [3, 26, 27]. Briefly, the contour of single MT segments was estimated within user-defined rectangular regions, and MT backbone coordinates were estimated by fitting a Guassian curve to each vertical line-scan of the region. Approximately 0.5µm interval of x-y coordinates were used to calculate MT curvature. The average Guassian curve amplitude of vertical line-scan at five highest curvature was expressed as MT AFU.

#### Manual MT tracking

To trace MT shapes in live cells and *in vitro*, we used protocols previously described in our laboratories [3, 4]. Briefly, in live cells, two representative regions of interest (ROIs) were selected and three representative single, clearly visible MTs were selected per ROI. Using ImageJ software (Fiji), x-y coordinate data from fluorescent images every 0.5 ± 0.05µm along 5-10 µm length of single clearly visible microtubules was collected in RFL-6 cells and *in vitro*. For MT Arbitrary Fluorescence Intensity (AFU), the fluorescent intensity of MT at each coordinate and adjacent background were recorded with circular selection which covers the entire width of the MT. The background intensity was subtracted from MT intensity at each coordinate and the average of MT fluorescence intensity per single MT was expressed as MT AFU.

### MT curvature estimation

Once x-y coordinates of MT contour were extracted semiautomatically or manually, the curvature (*k*) was calculated locally at each coordinate by taking three adjacent points and computing the change in the angle (*ϕ_k_*) over the average arc length of the two adjacent segments (Δ*S_k_*_-1_ and Δ*S_k_*) as we have previously described [3, 26]

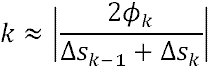

For the curvature distribution with five highest curvature, five highest curvature values from each MT were plotted.

### Fluorescence intensity analysis

The protocol to measure fluorescence intensity of GFP-UNC-45A on MT lattice was adapted from previously published work [56]. In order to measure GFP-UNC-45A signal at the same coordinate where MT curvature was measured, circular region of interest (ROI) covering the entire width of MT and the integrated density from the ROI was measured on the GFP-UNC-45A channel. The background intensity of GFP-UNC-45A channel nearby MT curvature measurement was subtracted from ROI intensity to remove the background noise signal. This background intensity values were also reported to ensure that the GFP-UNC-45A channel had homogeneous background. In cells, GFP mass AFU and p-S19 MLC mass AFU in RFL-6 cells were quantified as previously described [13].

### Statistical analysis

Results are reported as mean±s.d. of three or more independent experiments. Unless otherwise indicated, statistical significance of difference was assessed with an unpaired two-tailed Student’s *t*-test using Prism (V.4 GraphPad) and Excel (Microsoft). The level of significance was set at *P*<0.05.

## Data Availability

Data are available from the corresponding authors upon reasonable request.

## Supporting information

This article contains supporting information.

## Supporting information

Leged for Supporting Figures

## Acknowledgments

We thank Dr. Guillermo Marques (University of Minnesota Imaging Center) for assistance with image analysis.

## Author contributions

Conceptualization: A.H., M.B.; Methodology: A.H., B.C, D.O and M.B.; Software: A.H., B.C and M.B.; Validation: A.H., M.S., V.C., M.B.; Formal analysis: A.H., V.C., M.S., M.B.; Investigation: A.H., M.S., V.C. M.B.; Resources: M.B.; Data curation: A.H., M.B.; Writing: A.H., M.B.; Visualization: A.H.; Supervision: M.B.; Project administration: M.B.; Funding acquisition: D.O and M.B.

