## Supplementary figures and images for "The Microtubule Severing Protein UNC-45A Counteracts the Microtubule Straightening Effects of Taxol"

### Supporting Figure 1

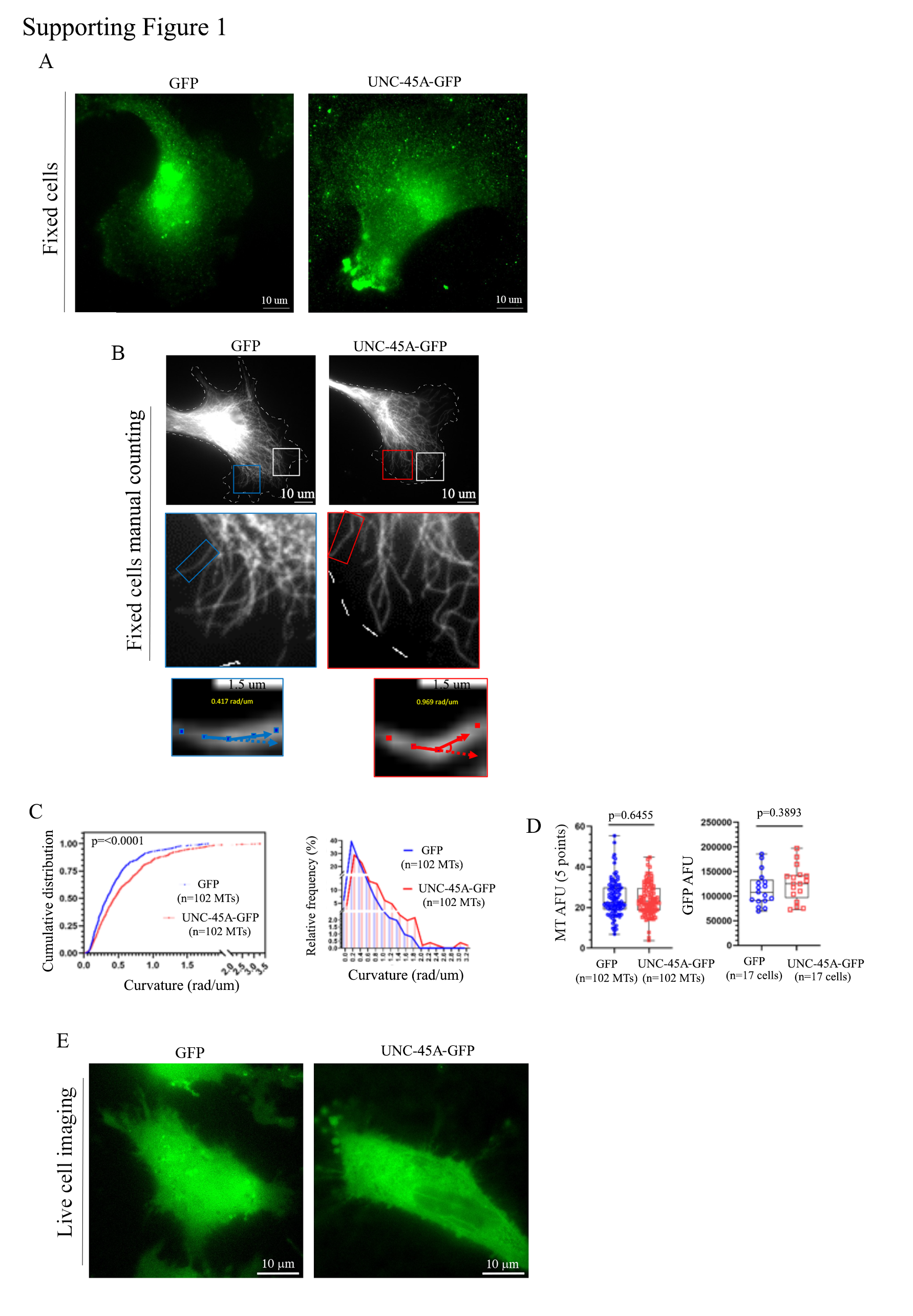

### Supporting Figure 2

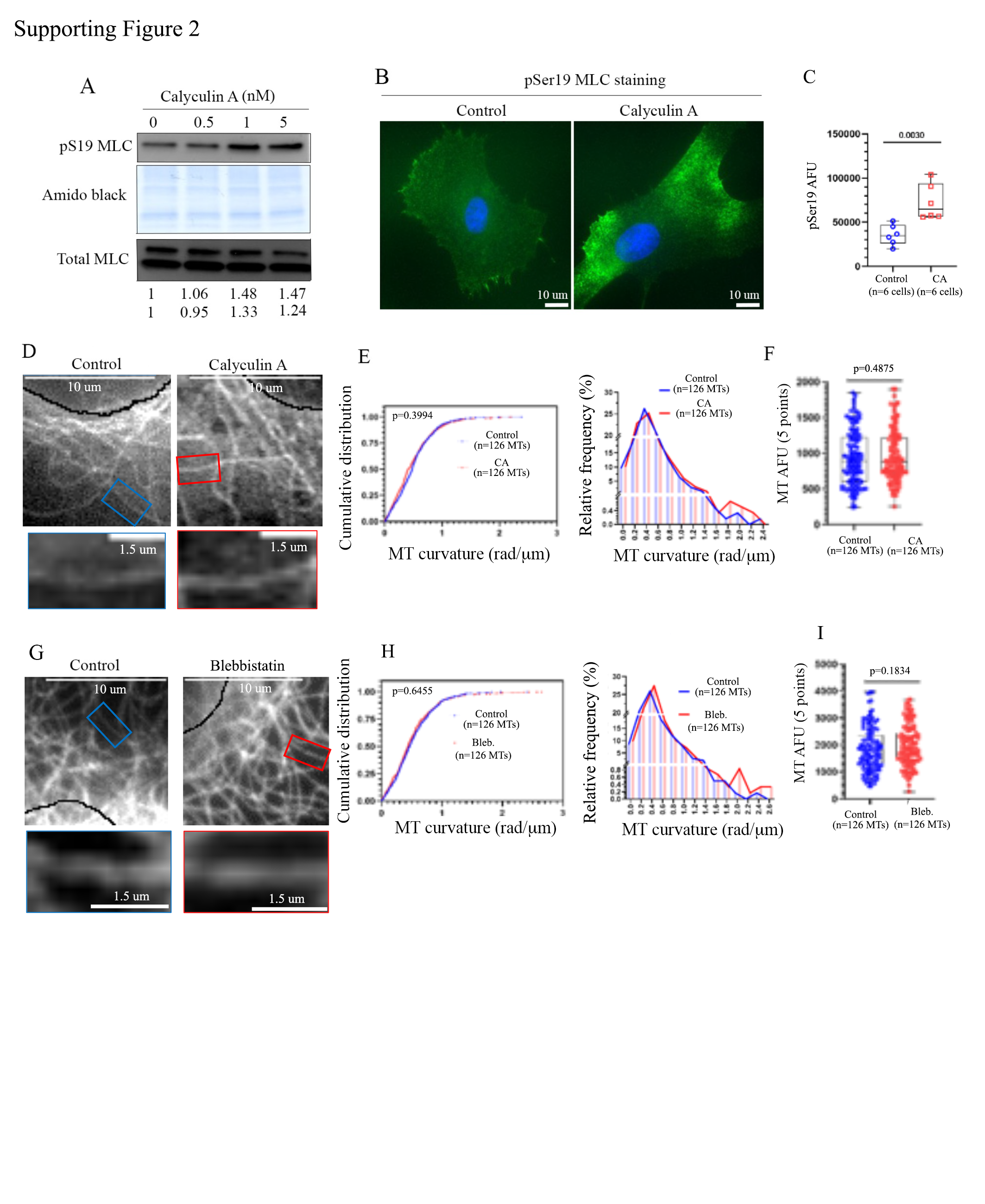

### Supporting Figure 3

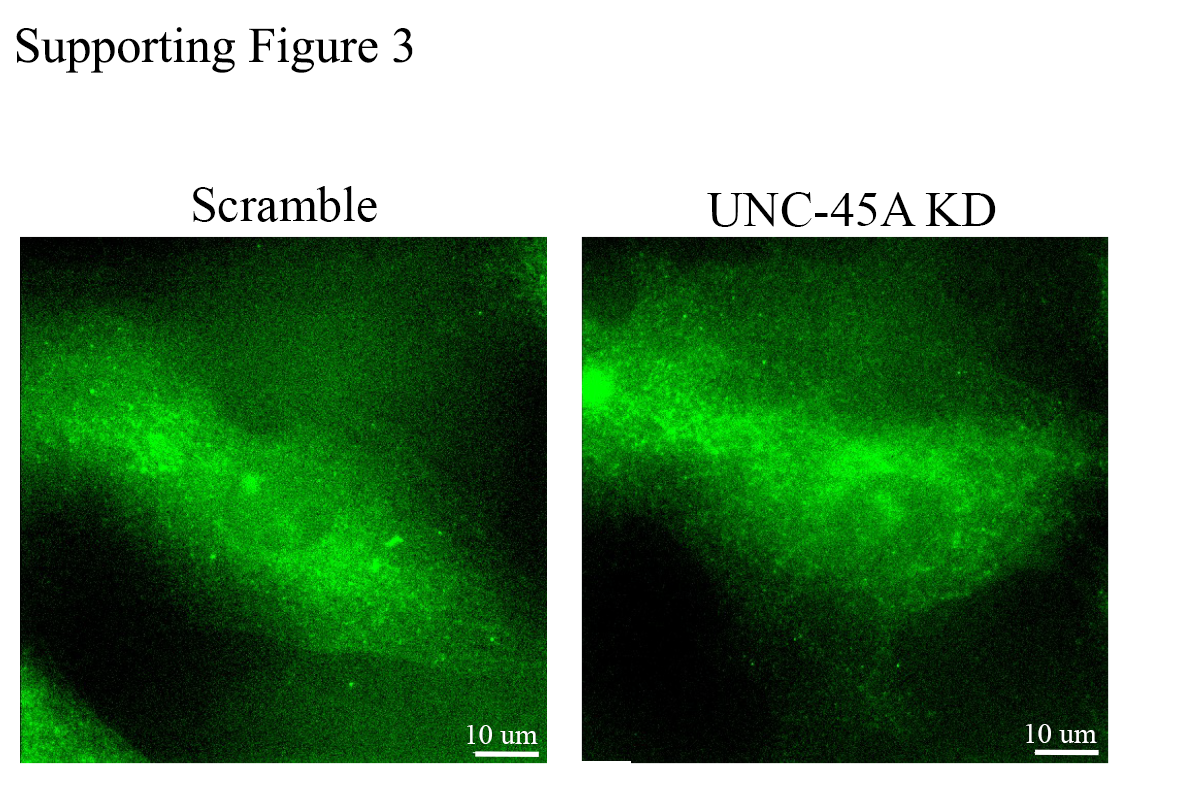

### Supporting Figure 4

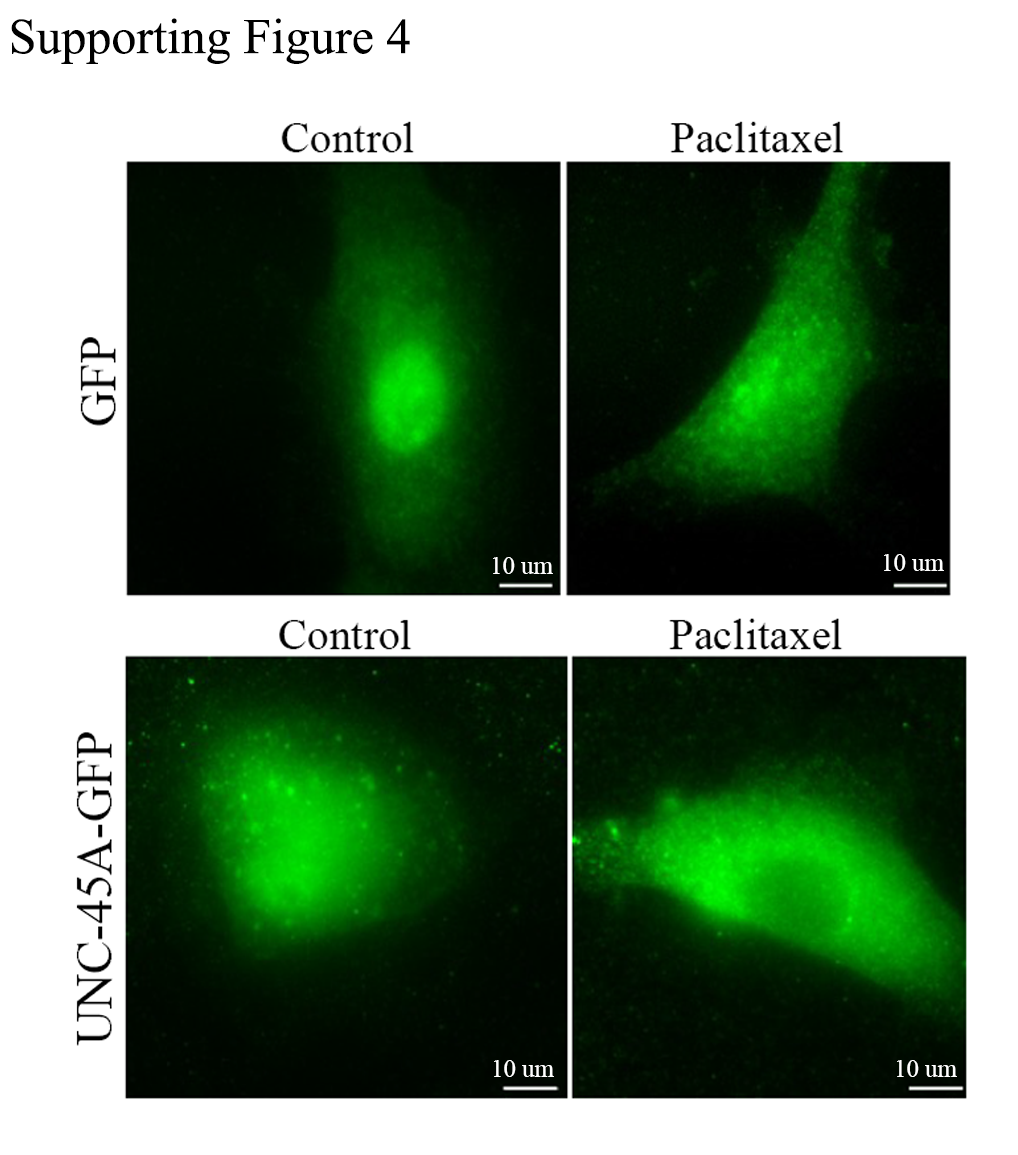
