## Supplementary material for "The Microtubule Severing Protein UNC-45A Counteracts the Microtubule Straightening Effects of Taxol": Leged for Supporting Figures

**Supporting Information**

**Supporting Figure 1. GFP and UNC-45A-GFP overexpressing cells and manual calculation of MT curvature in UNC-45A overexpressing cells. A**. Representative image of fixed GFP and UNC-45A-GFP overexpressing RFL-6 cells expressing similar amounts of GFP. **B**. *Top two panels,* representative images of peripheral MTs visualized via anti-alpha-tubulin staining in fixed GFP and UNC-45A-GFP overexpressing RFL-6 cells. The white dotted lines indicate cell edges. Images were taken with the same exposure time. Two representative regions of interest (ROIs) are shown. *Middle two panels,* close-up of one of representative areas shown in the top panels. *Bottom two panels,* representative magnified region of single MT shown in middle panels where 5 curvature values were obtained. For each MT, dots were placed every 0.5 µm for 2.5 µm long to record x-y coordinates. Yellow numbers indicate calculated curvature values at middle point using two adjacent points along MT. **C**. *Left,* cumulative distribution of MT curvature calculated using 5 curvature points per condition. The mean curvature values and standard deviations of GFP and UNC-45A-GFP were 0.430 rad/µm ± 0.339 and 0.583 rad/µm ± 0.477 respectively. *Right,* histogram of MT curvature distribution shown in C, *left*. **D**. *Left,* quantification of MT fluorescence intensity (arbitrary fluorescence units-AFU) along the length of the measured MTs. n= number of total MTs evaluated per condition. *Right,* quantification of GFP mass (arbitrary fluorescence units-AFU) in GFP and UNC-45A-GFPoverexpressing cells. **E.** Representative image of live GFP and UNC-45A-GFP overexpressing RFL-6 cells expressing similar amounts of GFP.

**Supporting Figure 2. Actomyosin contractility does not affect perinuclear MT curvature. A.** Western blot analysis of the levels of phosphorylated Ser-19 myosin light chain (p-S19 MLC) in RFL-6 cells treated with the indicated concentrations of calyculin A for 10 min. Amido black and total myosin light chain (MLC) were used as a loading control. Numbers represent the ratio between p-S19 MLC and amido black (*top*) and ratio between p-S19 MLC and total MLC (*bottom*) per condition. **B.** Representative images of RFL-6 cells treated with or without (control) calyculin A (1nM for 10 min), fixed and stained with p-S19 MLC (green) and DAPI (blue). **C.** Quantification of p-S19 MLC per condition expressed as AFU. Three different areas per cell were evaluated in six cells per condition. n=number of cells. **D.** *Top two panels,* representative images of perinuclear MTs visualized via anti-alpha-tubulin staining in fixed RFL-6 cells treated or not (control) with calyculin A (1nM for 10 min). Black lines indicate nuclear membrane. Images were taken using the same exposure time. Rectangles indicate the representative areas shown in the bottom panels. *Bottom two panels*, representative magnified region of single MT where 5 curvature values were obtained. **E.** *Left,* cumulative distribution of MT curvature calculated using the 5 curvature points per condition. The mean curvature values and standard deviations for control and calyculin A condition were 0.513 rad/µm ± 0.337 and 0.495 rad/µm ± 0.367 respectively. *Right,* histogram of MT curvature distribution shown in E, left. **F.** Quantification of MT fluorescence intensity (arbitrary fluorescence units-AFU) along the length of the measured MTs. n=number of total MTs evaluated per condition. Specifically, twenty one MTs per cell were evaluated in six cells per condition. **G.** *Top two panels,* representative images of perinuclear MTs visualized with alpha-tubulin staining in fixed RFL-6 cells treated or not (control) with blebbistatin (25$\mu$M for 1h). Black lines indicate nuclear membrane. All images were taken using the same exposure time. Rectangles indicate the representative areas shown in the bottom panels. *Bottom two panels*, representative magnified region of single MT where 5 curvature values were obtained. **H.** *Left,* cumulative distribution of MT curvature calculated using the 5 curvature points per condition. The mean curvature values and standard deviations for control and blebbistatin condition were 0.522 rad/µm ± 0.349 and 0.512 rad/µm ± 0.392 respectively. *Right,* histogram of MT curvature distribution shown in H, left. **I.** Quantification of MT fluorescence intensity (arbitrary fluorescence units-AFU) along the length of the measured MTs. n=number of total MTs evaluated per condition. Specifically, twenty one MTs per cell were evaluated in six control and blebbistatin treated cells.

**Supporting Figure 3. GFP expression in scramble and UNC-45A KD cells infected with a lentiviral plasmid expressing GFP.** A. Representative image of fixed scramble and UNC-45A KD RFL-6 cells expressing similar amounts of GFP.

**Supporting Figure 4. GFP and UNC-45A-GFP overexpressing cells treated or not with Paclitaxel.**

Representative image of fixed GFP and UNC-45A-GFP overexpressing RFL-6 cells treated or not (control) with paclitaxel expressing similar amounts of GFP.
